# Photobleaching step analysis for robust determination of protein complex stoichiometries

**DOI:** 10.1101/2020.08.26.268086

**Authors:** Johan Hummert, Klaus Yserentant, Theresa Fink, Jonas Euchner, Dirk-Peter Herten

## Abstract

The composition of cellular structures on the nanoscale is a key determinant of macroscopic functions in cell biology and beyond. Different fluorescence single-molecule techniques have proven ideally suited for measuring protein copy numbers of cellular structures in intact biological samples. Of these, photobleaching step analysis poses minimal demands on the microscope and its counting range has significantly improved with more sophisticated algorithms for step detection, albeit at an increasing computational cost. Here, we present a comprehensive framework for photobleaching step analysis, optimizing both data acquisition and analysis. To make full use of the potential of photobleaching step analysis, we evaluate various labelling strategies with respect to their molecular brightness and photostability. The developed analysis algorithm focuses on automation and computational efficiency. Moreover, we benchmark the framework with experimental data acquired on DNA origami labeled with defined fluorophore numbers to demonstrate counting of up to 35 fluorophores. Finally, we show the power of the combination of optimized trace acquisition and automated data analysis for robust protein counting by counting labelled nucleoporin 107 in nuclear pore complexes of intact U2OS cells. The successful in situ application promotes this framework as a new resource enabling cell biologists to robustly determine the stoichiometries of molecular assemblies at the single-molecule level in an automated fashion.

## Introduction

The fundamental functions of living cells are carried out by protein assemblies at the molecular level. Precise quantitative knowledge on the composition of these protein complexes in the cellular environment is crucial to further our understanding of their cellular functions. ^1^ In many cases these protein assemblies contain not only a variety of different components, but also several copies of each component, independent of whether the complex is a well-defined structure or a more variable cluster.^2^

To investigate the stoichiometry of a particular protein of interest in a molecular assembly, fluorescence microscopy offers several advantages. It is highly specific, live-cell compatible, single-molecule sensitive and therefore capable to resolve heterogeneities within ensembles in situ. In the last two decades, different fluorescence based molecular counting methods have been developed.^3^ Among them are methods relying on brightness calibration,^4^ counting of photobleaching steps,^5,6^ localization microscopy,^7–10^ or photon antibunching.^11^,^12^ To date, photobleaching step analysis (PBSA) and brightness estimation are most widely used in biological applications of molecular counting, due to their simplicity in data acquisition and the relatively straight-forward interpretation.^13^ PBSA has the advantage that no calibration measurements are necessary and that it is relatively robust to variations in molecular brightness.

While the idea of counting photobleaching steps is straight-forward, numerous approaches exist for data analysis.Often the number of steps is classified by visual inspection,^5,14,15^ which is not only time-consuming but also highly subjective. The exclusion of traces which cannot reliably be classified upon visual inspection will inevitably lead to a biased estimate, since traces with a higher number of fluorophores tend to exhibit a higher complexity. More reliable is the determination of the unitary step height by pairwise frequency analysis.^16^ Thereby, however, differences in step height over the field of view will broaden the measured emitter number distribution. Chung-Kennedy^6^ or median rank^15^ filters are often applied to photobleaching traces to improve step detection. Another noteworthy contribution to the technique is SONIC,^17^ which allows fast determination of fluorophore number means but cannot, for example, differentiate between unimodal or multimodal distributions. Recently molecular counting via photobleaching has attracted renewed attention due to the development of novel analysis modalities based on Bayesian statistics^18^,19 and machine learning.^20^

But these novel methods are demanding in terms of data quality, which in turn leads to new requirements regarding fluorophore properties. In the trade-off between signal-to-noise ratio (SNR) of the individual bleaching steps and the rate at which photobleaching occurs, bright and stable fluorophores are advantageous. Thus, buffer systems^21^,22 to achieve control over photophysical processes and to increase photostability could help to improve data quality. This motivates an investigation into which labelling approaches and buffer systems are most suited to generate data compatible with automated and robust photobleaching step analysis at an increased counting range. Additionally, Bayesian methods are computationally costly and therefore limit the number of photobleaching traces that can be processed in a given experiment. Therefore, we see the necessity for an approach to bridge the gap between simple methods such as visual inspection and the novel Bayesian methods. Lastly, the experimental validation with standard samples is often not the focus of theoretical methods development although calibration samples are readily available. ^23,24^

Here, we address these hurdles to make photobleaching step analysis a more robust and thoroughly validated tool in the biophysics toolbox. We describe a comprehensive framework for PBSA that provides guidelines for the choice of fluorophore label and acquisition conditions as well as a robust workflow for trace analysis. The trace analysis algorithm, which we named quickPBSA, is developed with a focus on automation, providing sufficient throughput to address the demands for in situ protein counting. We then validate the method with molecular counting experiments on DNA origami carrying defined label numbers. Finally, we show that quickPBSA, in conjunction with optimized trace data acquisition, is well suited to characterise protein structures in cells by determining the copy number of Nucleoporin 107 (NUP107) in the nuclear pore complex (NPC).

## Results

The reliability of automated photobleaching trace evaluation strongly depends on the quality of the input data, i.e. individual photobleaching traces. Historically, PBSA has mostly been performed using fluorescent proteins as labels.^13^ However, fluorescent proteins are typically less photostable than small organic fluorophores such as the carbopyronine ATTO 647N and are known to exhibit complex photophysics complicating trace interpretation.^25^ To identify fluorophores suited for generating photobleaching traces with high SNR, we evaluated the commonly used fluorescent proteins EGFP, mCherry and mNeonGreen with respect to their photostability. We further compared them to the two biocompatible small organic fluorophores tetramethylrhodamine (TMR) and silicon rhodamine (SiR) which can be conjugated to target proteins via self-labelling protein tags such as SNAP-tag or HaloTag.^26^ Finally, we tested to which degree photostabilizing buffers composed of reducing and oxidizing systems (ROXS) and an oxygen scavenger could be used to improve photostability and hence, trace quality.

Since the photostability of a fluorophore can strongly depend on its environment and on the applied measurement conditions, we performed systematic photobleaching measurements under comparable conditions with the three selected fluorescent proteins as well as with TMR and SiR substrates for both SNAP-tag and HaloTag. For this, we targeted each of the fluorescent proteins or protein tags to cellular structures via expression of fusion proteins in U2OS or COS-7 cell lines (Fig S1) and chemically fixed the cells prior to imaging. We then determined the photostability of each sample in phosphate buffered saline (PBS, pH 7.4) and three photostabilizing buffers each containing methyl viologen and ascorbic acid as ROXS and a oxygen scavenger system consisting of either glucose oxidase and catalase (GodCat),^27^ protocatechuate-3,4-dioxygenase (PCD)^28^ or sodium sulfite (NaSO_3_).^29^ While both, the GodCat and PCD systems have been extensively used for single-molecule fluorescence microscopy, NaSO_3_ has only recently been used as additive in photo-switching buffers for single-molecule localization microscopy.

In contrast to the photostability, the photophysical properties of fluorescent proteins and small organic fluorophores are well studied and can be readily compared based on reports in the literature (Table S1). For protein tag conjugated fluorophores, a strong influence of the protein tag on the fluorophore substrate was recently reported and has to be taken into account when comparing the brightness of different fluorophores after conjugation to protein tags.^30^ To identify promising labelling strategies suited for PBSA of cellular targets, we then compared the measured photostabilities as well as the nominal brightness of each fluorophore with ATTO 647N, which is a well suited fluorophore for in vitro measurements due to its high molecular brightness and excellent photostability.

Under the high intensity illumination in the widefield epifluorescence microscope we observed varying photobleaching decay patterns for the tested fluorophores (Fig S1). We therefore decided to use the time to reach half maximum *t*_1*/*2_ as a model-independent metric to compare conditions and fluorophores (Fig 1a). Across all conditions, we observed substantial differences in *t*_1*/*2_ covering three orders of magnitude (0.5 - 200 s). Among the tested fluorophores, fluorescent proteins were less photostable than organic fluorophores and far-red organic fluorophores exhibited the highest photostability. Differences between fluorophores were less pronounced in conventional buffers without photostabilizing additives (at most 8-fold difference across conditions). Addition of ROXS and oxygen scavengers resulted in strongly increased photostability for organic fluorophores, particularly for the red-absorbing fluorophores ATTO 647N and SiR (Fig 1a,b). For ATTO 647N, photostability was improved 2-fold with ROXS buffer supplemented with NaSO_3_ as oxygen scavenger. Replacing NaSO_3_ with the enzymatic oxygen scavengers GodCat or PCD resulted in further improvements of 10 and 20-fold respectively. Similar trends were observed for both TMR and SiR conjugated to SNAP-tag or HaloTag. Here, SiR conjugated to SNAP-tag in ROXS PCD showed the highest stability with a *t*_1*/*2_ of 120 s (20-fold improvement over PBS), a 60-fold improvement over EGFP, but still 50% lower stability compared to ATTO 647N in the same buffer. In contrast, a decrease in photostability for EGFP and mCherry in the methyl viologen and ascorbic-acid-based ROXS buffer was observed for all three oxygen scavenging systems, possibly due to pH changes of the buffers during imaging.^31^ Interestingly, no such decrease in photostability was observed for mNeonGreen, which could be due to improved stability ofthe protein structure of mNeonGreen.^32^

**Figure 1:**
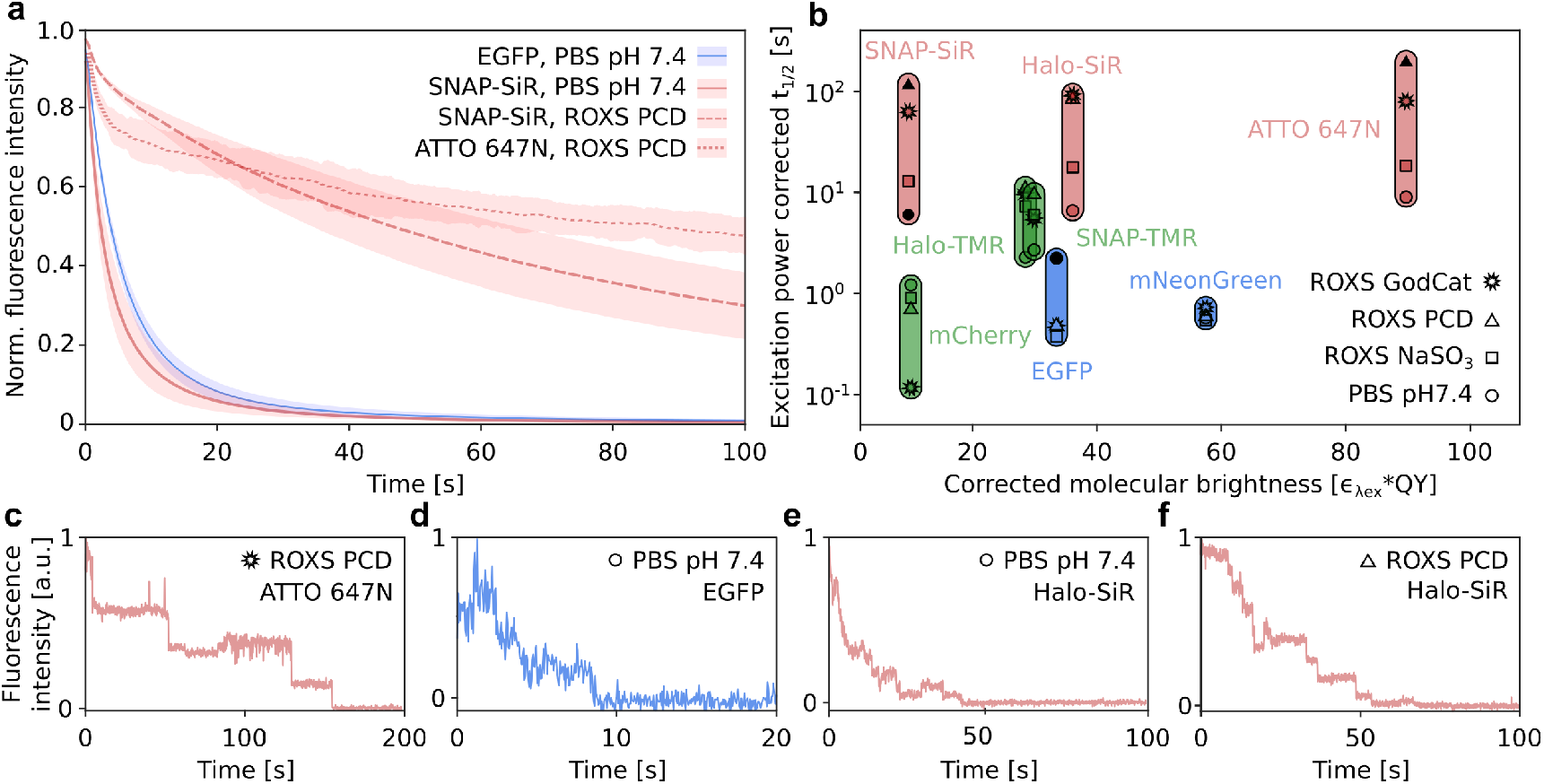
Improved fluorophore and image acquisition buffer selection for PBSA. Different fluorescent proteins and substrates for self-labelling protein tags were evaluated with respect to their photostability, brightness and propensity for photo-induced blinking. **a**, Normalized photobleaching traces for indicated fluorophore-buffer combinations. Mean (line) and standard deviation (shaded region) of 7-10 measurements from 2 independent experiments per condition. **b**, Excitation power density corrected photostability vs. excitation wavelength corrected brightness for individual fluorophore – buffer combinations. Symbols indicate mean photostability under respective condition. Conditions shown in a and c-f highlighted as bold symbols. Colours indicate excitation wavelengths used (488 nm – blue, 561 nm green, 640 nm – red). See Fig S2 for full dataset. **c**, Representative intensity transients from indicated conditions. Traces were normalized to their respective maximum and minimum values.

In addition to increasing the photostability of organic fluorophores, ROXS buffers were also reported to reduce emission intensity fluctuations on the ms timescale.^22^ Such blinking was frequently observed for fluorescent proteins in all buffers tested (see e.g. EGFP, Fig 1d), while it could be efficiently suppressed for ATTO 647N in ROXS GodCat buffer (Fig 1c). To identify buffer compositions able to efficiently suppress blinking for protein tag conjugated fluorophores, we visually inspected photobleaching traces from the individual conditions. For all organic dyes we found that ROXS buffers containing any of the three tested oxygen scavengers reduced blinking (Fig 1e,f), however not to the degree observed for ATTO 647N. In contrast to small organic fluorophores, we observed no substantial change in blinking behaviour for fluorescent proteins in any of the test buffer compositions. Based on these observations, we hypothesized that using self-labelling protein tags in combination with long-wavelength fluorophores and ROXS buffers supplemented with enzymatic oxygen scavenger systems, bears the potential to improve trace quality and thereby to extend the counting range of PBSA-based protein counting.

Turning from the data acquisition to the data analysis, we set out to develop a routine capable of estimating fluorophore number distributions directly from the experimental data (i.e. image stacks) with minimal user input. To that end quickPBSA includes modules for automated trace extraction from raw time-lapse image stacks and filtering as well as a new algorithm for trace interpretation. The underlying principle of the trace analysis algorithm is to perform a preliminary step detection for each trace and then refine the results iteratively. The final refinement step makes use of a Bayesian posterior defined by Pressé and coworkers,^18^ thus incorporating prior knowledge about the photobleaching process. The full data analysis workflow is comprised of four major parts (Fig 2a) which are described in the following:

**Figure 2:**
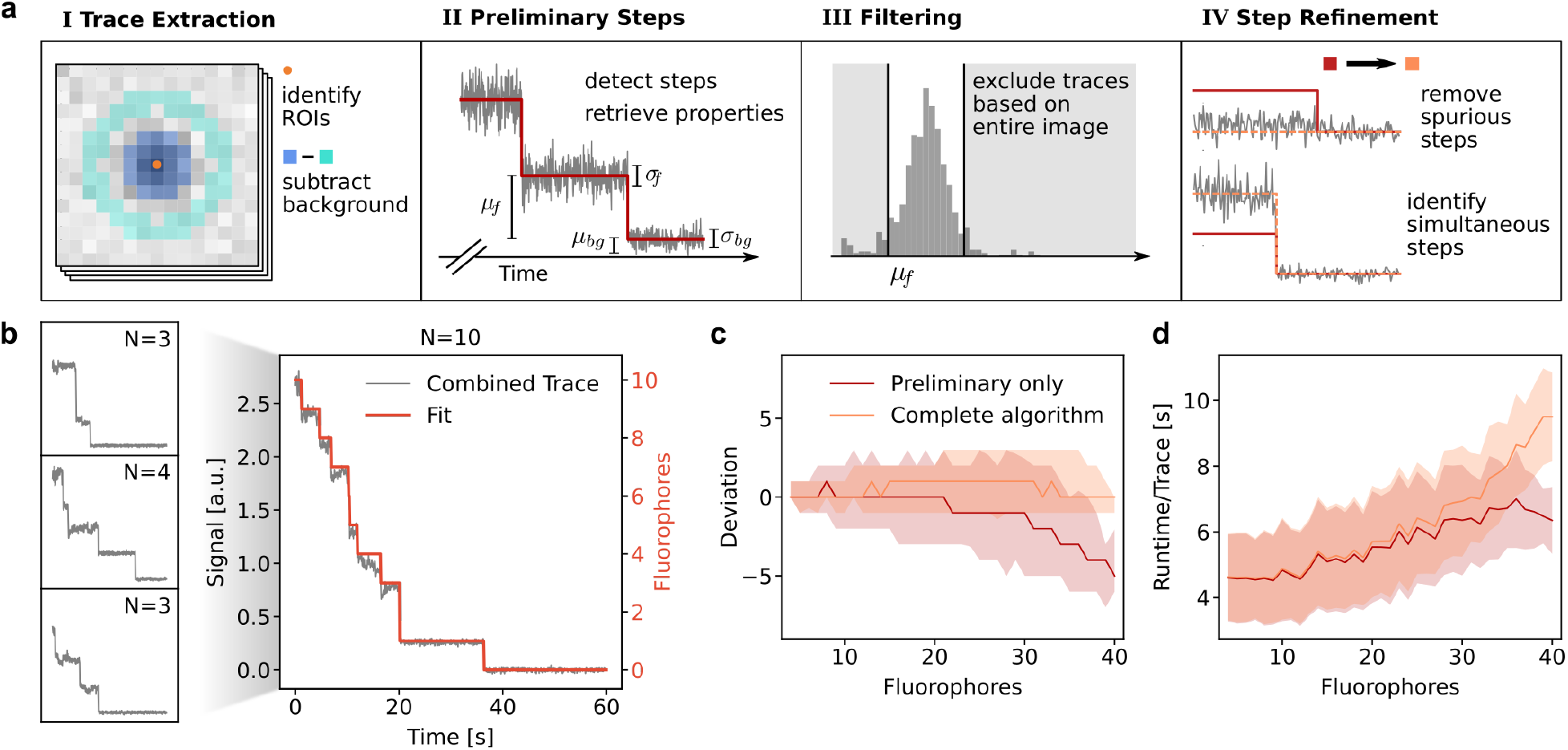
Algorithm and benchmarking. **a**, Main parts of the analysis framework as detailed in the text. **b**, Semi-synthetic traces for benchmarking are generated by combining manually classified traces. **c**, Deviation from ground truth for the benchmark with semi-synthetic traces. Median value (line) and the 5^th^ and 95^th^ percentiles (shaded region) are shown. **d**, Runtime benchmark result. Mean (line) and standard deviation (shaded region) are shown.

**I**.The identification of regions of interest (ROIs) and the automated extraction of photo-bleaching traces from image stacks. Here, the ROIs can be provided as pixel coordinates or via a segmentation mask image. As the photobleaching trace is extracted from the ROI, a ring-shaped region with variable offset from the ROI is used to extract a background bleaching trace (Fig 2a). Other ROIs are automatically excluded from the background region. Especially for measurements in cells we found that background bleaching due to autofluorescence and out-of-focus fluorescence occurred on similar timescales as the fluorophore bleaching. Therefore, background subtraction proved to be essential to recover traces with discernable photobleaching steps. Additionally, background subtraction also facilitates identifying and excluding ROIs that are not fully bleached at the end of the measurement.

**II**.After trace extraction, a preliminary step detection is performed on all extracted traces. This is accomplished by successively placing steps at positions where large changes in the mean intensity occur and evaluating each added step using the Schwarz Information Criterion (SIC) as first demonstrated by Kalafut and Visscher.^33^ In our implementation of this algorithm, a user-defined threshold below which changes in the mean intensity are ignored, reduces the detection of spurious steps rendering the preliminary step detection more robust (see SI).

**III** Traces are excluded based on the result of the preliminary step detection. The model selection in part IV critically depends on the correct detection of the last and the penultimate bleaching steps in each trace since the period where only one fluorophore is active is used to retrieve the properties of an individual fluorophore. Therefore, traces which are not fully bleached at the end of an acquisition or where the last step putatively corresponds to a double bleaching event need to be excluded. Assuming that the last two steps are correctly identified in a majority of traces from the image stack, the distribution of single fluorophore signal means across all traces can be used to filter out traces. By excluding traces where the single fluorophore signal or the background signal are out of bounds, we use information from the entire image stack to identify traces where the last two steps have been correctly located.

**IV** Ultimately, the result is refined by evaluating the entire trace according to the full marginal posterior from Pressé and coworkers.^18^ This posterior incorporates the possibility of simultaneous bleaching events as well as a penalty for too many bleaching events and thus is a far more content-aware evaluation of step placement than the information criterion used in the preliminary step detection (see SI). The algorithm iteratively minimizes the negative logarithm of the posterior, starting from the result of the preliminary step placement with all steps considered to be single events. A detailed flowchart of the refinement is shown in Fig S3. The iterative procedure is:

1. Find steps with multiple occupancy by trying all possible combinations of step positions for double steps. For higher occupancy, only the locations from the last iteration are considered, meaning we consider triple steps only where double steps yielded an improved posterior. All occupancies are reset to one in every iteration so that, for instance, [2,2,2] can be replaced by [1,3,2]. The result of this process is accepted and considered the new optimum, if a lower value of -log(P) is found at any point.
2. Remove the least significant step found during preliminary step detection. The final two steps in a trace are always kept in place, since they are required for posterior calculation. If there are no steps left to remove proceed with step 3, else reset all steps to single occupancy and return to step 1.
3. If no steps could be removed to yield an improved posterior, i.e. the current optimum contains the same number of steps as the preliminary result, the algorithm proceeds to add single occupancy steps. This is accomplished by calculating -log(P) for additional positive or negative steps at all positions before the penultimate step. Repeat step 3 until the last two added steps yielded no improved posterior or a specified maximum number of added steps is reached.
4. Return the step/event combination with the minimal value for -log(P) found at any point in 1-3.

In part IV of the workflow, the evaluation of simultaneous step arrangements is computationally most expensive. Especially for traces with many steps in the preliminary result, the number of possible combinations is excessive. We therefore implemented several strategies to reduce the computational cost at this point, as detailed in the documentation of the software package (see Methods).

We benchmarked the quickPBSA trace analysis algorithm using semi-synthetic data generated from experimental data where the ground truth could be obtained by manual inspection of traces. For this, we acquired experimental data from an in vitro sample with few fluorophores per diffraction-limited spot, namely immobilized DNA oligonucleotides labeled with four ATTO 647N fluorophores (see Methods for details). We selected traces where the quickPBSA result could be confirmed by visual inspection, obtaining a set of traces with known ground truth. We then generated increasingly complex semi-synthetic traces with known ground by combining several of these traces (Fig 2b). Using this approach, bench-marking traces with fluorophore numbers up to 40 were generated and used to evaluate the accuracy of the quickPBSA trace analysis (Fig 2c).

For each fluorophore number, approximately 100 semi-synthetic traces with 3,000 time-points per trace were included in the analysis (Fig S3b). We observed that even for these hand-picked traces, the result of the preliminary step detection started to deviate systematically from the ground truth for fluorophore numbers beyond 20, most likely due to missed bleaching events which occurred in close temporal succession. Although the trace refinement in the complete quickPBSA algorithm favours removing steps over adding them, the quickPBSA result did not exhibit a systematic bias. The median estimated fluorophore numbers never deviated by more than one fluorophore from the expected value (Fig 2c).

We also used the semi-synthetic traces to benchmark the runtime of the analysis in dependence on the number of fluorophores (Fig 2c). We observed that for up to 20 fluorophores, the runtime was dominated by the preliminary step detection. For higher fluorophore numbers the runtime increase due to the result refinement (IV) started to contribute and increased further for higher fluorophore numbers. Nonetheless, the mean runtime remained below 10 s per 3,000 frame trace for the entire benchmarking dataset containing traces with up to 40 fluorophores. To compare this runtime with a state-of-the-art Bayesian method, we analysed a subset of the benchmarking dataset with a previously published software package.^18,34^.The subset consisted of 40 traces, containing two initial ground truth traces each (i.e. 2-8 fluorophores). For this subset of the benchmarking data, we recorded an average runtime of 75*±*15 min per trace. Since this result is close to the specified runtime of 1 s/frame,^34^ we expect that at most a two-fold improvement can be achieved by parameter tuning. Thus, using the quickPBSA algorithm, we were able to analyse the dataset with an approximately 1,000-fold lower computation time. The mean analysis times for all datasets included in this publication are below 3 minutes per trace even for traces with up to 15,000 data points (Table 1).

**Table 1:** Mean and standard deviations extracted from Gaussian modelling of measured emitter number distributions from DNA origami experiments. Standard errors in brackets as extracted from least squares fitting. The expected values are calculated assuming a 70% labelling efficiency. *Measurement on alternative microscope setup with sCMOS detector. Std. Dev.: Standard deviation.

| Sample | Mean | Mean expected | Std. Dev. | Std. Dev. expected | # Traces | Runtime/Trace [s] |
| --- | --- | --- | --- | --- | --- | --- |
| R09 | 6.7 (0.1) | 6.3 | 1.9 (0.1) | 1.4 | 197 | 25 |
| R20 | 14.2 (0.2) | 14.0 | 6.1 (0.2) | 2.1 | 636 | 53 |
| R35 | 22.6 (0.5) | 24.5 | 8.6 (0.5) | 2.7 | 499 | 168 |
| R09* | 6.0 (0.1) | 6.3 | 2.2 (0.1) | 1.4 | 1,667 | 19 |
| Y09 | 7.9 (0.1) | 6.3 | 1.6 (0.1) | 1.4 | 853 | 88 |

To fully validate the developed framework with experimental data, we used DNA origami carrying a well-defined number of fluorophores. DNA origami with 9, 20, and 35 binding sites for labelling strands carrying a single ATTO 647N fluorophore (R09, R20, and R35) were sparsely immobilized on coverslips to ensure that stochastically overlapping origami structures did not significantly influence the measurement (Fig 3a left, see Methods). Photobleaching traces from individual DNA origami structures were then extracted using the trace extraction module described above (see Methods). Since the extracted traces exhibited only weak background bleaching, the background subtraction step for this sample mainly removed a constant offset caused by excitation bleed-through and read-out noise (Fig 3a center). Processing the background-corrected traces using the full quickPBSA algorithm resulted in good agreement between intensity and predicted fluorophore numbers over time (Fig 3a right). The obtained fluorophore number distributions were symmetrical indicating no systematic deviation and an unbiased measurement error (Fig 3b). The means obtained from fitting a normal distribution to the quickPBSA fluorophore number estimates were in excellent agreement with the expected values for a labelling efficiency of 70% (see Methods). In contrast, the means of distributions from only the preliminary step detection exhibited a clear underestimation for origami with 20 and 35 binding sites (Fig 3c), confirming the initial benchmarking results with semi-synthetic traces.

**Figure 3:**
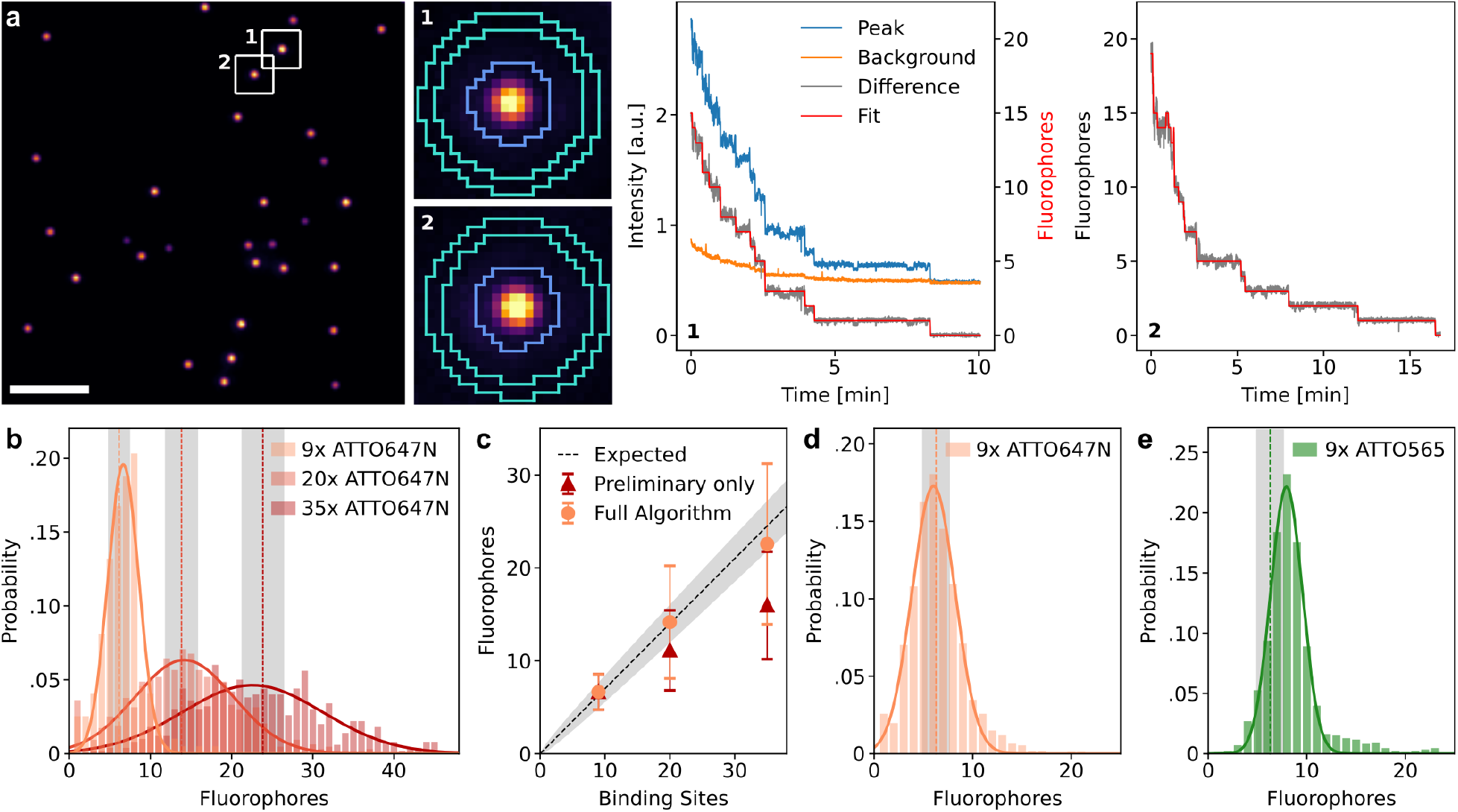
Validation with DNA origami samples **a**, Representative image and traces from the origami experiment with 20 binding sites for ATTO 647N. Scale bar: 10 *µ*m. **b**, Fluorophore number distributions for origami with 9, 20, and 35 binding sites. The histograms are modeled with a Gaussian to extract means and standard deviations (results in table 1). Vertical dashed lines and areas shaded in grey indicate the expected mean and standard deviation obtained from binomial distributions. **c**, Fit results from b compared to the expected mean of the label number distribution, which is a binomial distribution with a labelling efficiency of 70%. Error bars and shaded region show the standard deviation. **d**, Measured label number distribution of origami with 9 binding sites for ATTO 647N on a different microscope setup with a larger field of view and sCMOS detector. Vertical line and shaded region indicates expected mean and standard deviation. **e**, Measured label number distribution for origami with 9 binding sites for ATTO 565. Vertical line and shaded region indicates expected mean and standard deviation.

A full comparison of all parameters from the fits and a comparison to the mean and standard deviation of a binomial distribution with the known number of binding sites and the expected labelling efficiency is shown in Table 1. We observed that the measured distributions broadened with increasing fluorophore number, stronger than would be expected from the binomial distribution of label numbers alone. For instance, while broadening of the measured data was negligible for R09 origami, the standard deviation increased by a factor of 3 for R20 origami (Table 1). This suggests that experimental data contains additional sources of uncertainty which are not fully reproduced using semi-synthetic data and therefore highlights the importance of additional benchmarking with standardized samples.

To validate that the quickPBSA algorithm performed robustly upon variation of experimental parameters, we performed additional measurements with the R09 origami samples on a different widefield microscope setup with homogeneous illumination power and a sCMOS instead of an emCCD camera for detection (see methods for details). Also in this experiment the expected mean and width of the fluorophore number distribution were well reproduced (Fig 3d, Table 1). A large field of view is advantageous for the acquisition of photobleaching data since overall measurement time is decreased and the potential impact of degrading buffer performance can be minimized.

We furthermore explored the sensitivity of the quickPBSA algorithm to fluorophore properties by measuring a fluorophore number distribution for origami with 9 binding sites labelled with ATTO 565 (Y09). Here, the measured distribution showed a peak at 7.9 fluorophores, significantly above the expected mean fluorophore number of 6.3 (Table 1). A likely explanation for this deviation is that ATTO 565 exhibited two distinct brightness states as frequently observed in individual photobleaching traces (Fig S4). If the last photobleaching step occurs from a lower brightness state in a significant number of traces, the mean signal of a single fluorophore is underestimated for these traces, leading to an overestimation of the total number of fluorophores. This again highlights how important label selection is in photobleaching experiments, even for organic fluorophores. Additionally, taking the complex photochemical behaviour into account can allow to extend the counting range of photobleaching analysis even further.^35^

While DNA origami samples are ideally suited for determining the accessible counting range of a novel method, the application to biological targets in situ, i.e. within the complex cellular environment is subject to additional challenges that are not captured in such simplified in vitro experiments. Background (auto)fluorescence, density of structures and biological variation cannot readily be controlled in a biological sample and will impact data quality.

To assess how the quickPBSA framework coped with a more complex in situ sample, we decided to determine the number of NUP107 protein copies contained in individual nuclear pore complexes. To minimize the influence of protein expression and labelling efficiency, we used a genome-edited U2OS cell line expressing NUP107-SNAP from its native genomic context.^36^ Labelling of SNAP-tag conjugated NUP107 was performed with the corresponding silicon rhodamine derivate BG-SiR.

From epifluorescence images of chemically fixed and labeled cells, it is immediately evident that fluorescent background is much more pronounced in situ than in the origami experiments described above (Fig 4a). Additionally, the high density of NPCs resulted in regions where it was no longer possible to identify individual NPCs. We therefore excluded regions with overlapping NPCs and only considered sufficiently isolated and diffraction limited structures for further analysis. To this end, all pre-localized spots closer than a given threshold and spots with a width larger than the diffraction limit were excluded before trace extraction (Fig S5). Despite this pre-filtering, raw traces from individual ROIs did not exhibit clear bleaching steps and the decay in the background region occurred on a similar timescale as the fluorescence signal of the ROI (Fig 4b). Background fluorescence can therefore be mainly attributed to out-of-focus fluorescence rather than autofluorescence. After subtraction of the background signal, photobleaching steps could be observed towards the end of photobleaching traces (Fig 4b). Despite the fact that the obtained traces had a substantially lower SNR than the previously successfully evaluated traces recorded using ATTO 647 as fluorophore, we subjected the extracted traces to analysis with the quickPBSA algorithm (Fig 4b and 4c). The resulting fluorophore number distribution cumulated from 3 independent experiments (Fig S5) was well described by a normal distribution with a mean of 20.6*±*0.2 fluorophores per NPC and a standard deviation of 8.4*±*0.2 (Fig 4d). The width of the distribution is comparable to that of the distribution obtained from R35 DNA origami, indicating that background subtraction and spot pre-filtering successfully reduced the complexity of obtained traces and did not result in reduced precision. NUP107 has been reported previously to be present in NPCs at 32 copies per pore.^37^,38 Based on the mean fluorophore number of 20.6*±*0.2 per NPC, this translates into a labelling efficiency of 64% for SNAP-tag labelling with BG-SiR which is in excellent agreement with recent reports^24^ indicating that quickPBSA is able to correctly measure fluorophore numbers even for less bright fluorophore labels and in the complex environment of a eukaryotic cell.

**Figure 4:**
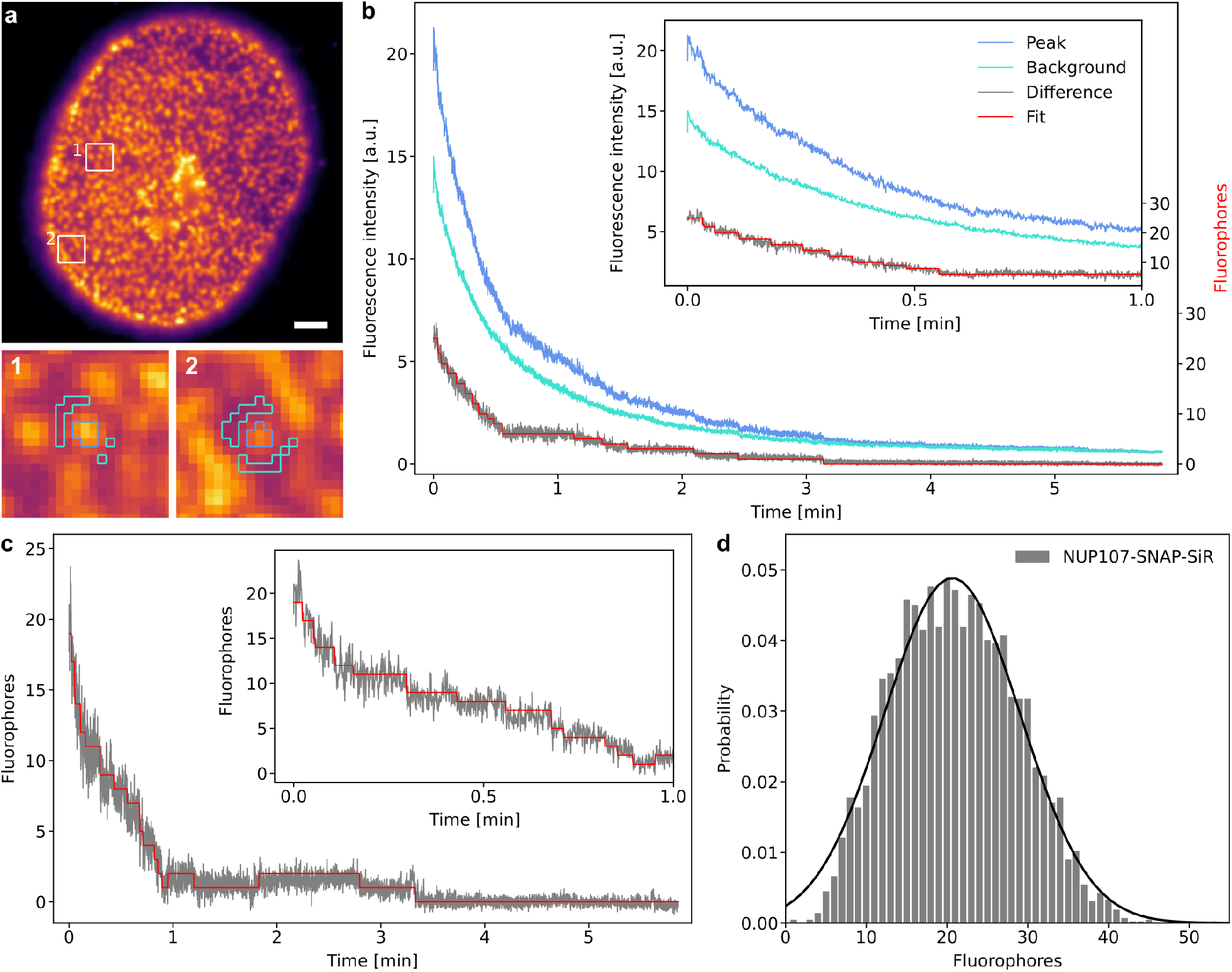
Protein counting of NUP107 in U2OS cells. **a**, Representative image of U2OS cell stably expressing NUP107-SNAP-tag labeled with BG-SiR. Scale bar: 5 *µ*m. **b**, Traces extracted from the segmented ROI and background regions and evaluated difference trace from example trace b. **c**, Evaluated background-corrected trace extracted from ROI c (scaled to overlap). **d**, Measured fluorophore number distribution and Gaussian model fit (black line, mean 20.6*±*0.2). 6,227 traces from 43 cells, 3 independent experiments.

## Discussion

The presented framework for photobleaching step analysis offers a robust, fast, and well-validated approach for molecular counting in situ. The evaluation of various fluorophores along with different buffer conditions in respect to the precision of photobleaching step analysis may serve as practical guide for robust counting of proteins and other biomolecules. Additionally, it may serve as blue print for extended screening of other fluorescent labels and experimental conditions. The high photostability of organic fluorophores imaged in ROXS buffer with enzymatic oxygen removal enabled generating high-quality input data for automated trace interpretation. Despite the lower brightness compared to its HaloTag equivalent, the excellent photostability of SiR conjugated to SNAP-tag in ROXS/PCD buffer was found to be suitable for PBSA protein counting. Clearly, photostability and brightness also strongly depend on the specific environment, as especially evident by the effect of protein tag conjugation on SiR (table S1). Therefore, evaluating fluorophore properties should always be considered when using alternative labelling approaches. Fluorophores with improved molecular brightness and photostability^39^,40 as well as recently reported self-healing fluorophores^41^,42 might allow to extend the accessible counting range of photobleaching step analysis. Approaches to improve the SNR during image acquisition such as confocalized detection or single-plane illumination could help to improve trace quality and thereby further extend the accessible counting range of quickPBSA.^43^

On the analysis side we found it beneficial to make use of information from the entire field of view during trace selection, combining features from pairwise frequency methods with features from Bayesian approaches. In this spirit, the methodology for trace analysis is a combination and extension of two previous approaches to photobleaching step analysis. The combined method has only few user-defined parameters simplifying automation and improving robustness. The high degree of automation together with the more than 100-fold improved computational efficiency of the combined method provide a significantly increased throughput required for biological applications. We believe that the developed method of testing algorithms with semi-synthetic data will be highly useful, not only for benchmarking other PBSA algorithms, but also to generate training data for machine learning based approaches.^20^

Using ATTO 647N-labelled structures with a known stoichiometry in vitro, we showed that quickPBSA yields highly accurate (*<*10% deviation across all samples) estimates of mean fluorophore numbers for structures containing up to 35 fluorophores. We furthermore demonstrated the robustness of the quickPBSA workflow by successfully analysing data acquired on different experimental setups. We also demonstrated that the complex photo-chemical behaviour of fluorophores can skew fluorophore number estimates highlighting the importance of careful fluorophore characterisation prior to experiments.

To show that quickPBSA performs well in biological applications, we used quickPBSA to determine the number of NUP107 protein copies in nuclear pore complexes of U2OS cells. At this point, the background subtraction and trace filtering modules of quickPBSA proved crucial to obtain traces from complex samples. Factoring in the expected labelling efficiency for SNAP-tag labelling, the previously reported number of 32 protein copies was well reproduced. This constitutes, to our knowledge, the highest stoichiometry successfully measured with photobleaching step analysis in a biological sample so far and demonstrates the robustness of the outlined approach in biological samples.

Future developments of alternative algorithms for PBSA to further improve the precision of fluorophore counting, such as novel Bayesian approaches,^35^ will be of high interest to move beyond measuring mean complex stoichiometries and towards characterizing stoichiometry distributions across ensembles of individual complexes. At this point, however, advances in data analysis will need to go hand in hand with the development of novel labelling schemes with improved labelling efficiency to reduce the variation in fluorophore numbers caused merely by incomplete labelling of target proteins.

Overall, the combination of improved data acquisition and the novel analysis routines contained in the quickPSBA framework provide a reliable way to determine protein stoichiometries in cellulo and will enable the use of automated PBSA as routine tool for cell biology in future applications.

## Methods

### Preparation of DNA *in vitro* samples

Custom brightness DNA origami with 9,20 or 35 nominal binding sites labeled with ATTO 647N or ATTO 565 at approximate labelling efficiencies of 70% (Gattaquant DNA Nanotechnologies, Germany) were dissolved in 0.5xTBE buffer supplemented with 11 mM MgCl_2_ and stored at -20°C until use. DNA oligonucleotides labeled with 4 ATTO 647N (tetraATTO 647N) as previously described^17^(biomers.net, Germany) were dissolved in phosphate buffered saline (PBS, 10 mM phosphate buffer, 2.7 mM KCl, 137 mM NaCl, pH 7.4, Sigma Aldrich, Germany) and stored at -20°C until use. Both, DNA origami and DNA oligonucleotides were immobilized in 8-well LabTek (Nunc/Thermo Fisher, US) chambered coverslips via biotin-streptavidin linkage as previously described.^44^ Prepared samples were kept in PBS (DNA oligonucleotides) or PBS supplemented with 20 mM MgCl_2_ (DNA origami) unless stated otherwise.

### Cell Culture

COS-7, HeLa and U2OS cells were cultured in Dulbecco’s modified eagle medium (DMEM) supplemented with GlutaMax and 1 mM sodium pyruvate (all Gibco Technologies, US). Cells were grown at 37°C, 5% CO2 in humidified atmosphere and routinely subcultured every 3 days or upon reaching 80% confluency. Cultures were kept for up to 30 passages. For widefield imaging, cells were seeded into 8-well LabTek chambered coverslips. Prior to seeding cells, LabTeks coverslips were cleaned with 0.1 M hydrofluoric acid for 2×30 s and extensively washed with PBS. Transfection of COS-7 cells was performed with TransIT-X2 transfection reagent (Mirus Bio, US) according to manufacturer’s instructions 24 hours after cell seeding and at least 22 hours before fixation. Fixation of cells was performed with 3.7% (w/v) pre-warmed paraformaldehyde (PFA, EM Grade, Electron Microscopy Sciences, US) freshly diluted in PBS for 20 minutes at room temperature unless stated otherwise. U2OS cells expressing NUP107-SNAP-tag were pre-fixed with 2.4% PFA (w/v) in PBS for 30 s, permeabilized with 0.4% (w/v) Triton X-100 in PBS for 3 min and then fixed with 2.4% PFA in PBS for 30 min. All samples were washed repeatedly with PBS after fixation and imaged directly after or kept in PBS at 4°C until being imaged.

### Preparation of cells expressing SNAP-tag or HaloTag

HeLa cells stably expressing GlnA-HaloTag ^45^ were a kind gift of Florian Salopiata (DKFZ Heidelberg). U2OS cells stably expressing NUP107-SNAP-tag were a kind gift of Jan Ellenberg (EMBL Heidelberg). ^46^ Both cell lines were labeled with corresponding tag substrates directly prior to fixation. GlnA-HaloTag expressing cells were labeled with tetramethyl rhodamine (TMR) HaloTag ligand (HTL-TMR) or silicon rhodamine HaloTag ligand (SiR-HTL) at 100 nM in growth medium for 120 min at 37°C. NUP107-SNAP-tag expressing cells were labeled with benzylguanine-functionalised TMR or SiR (BG-TMR/SiR) at 200 nM in growth medium for 120 min at 37°C. After labelling, cells were washed repeatedly with growth medium and fixed as described above.

### Imaging buffers

Samples were imaged either in PBS or in buffers containing different reducing and oxidizing systems (ROXS). ROXS buffers were prepared from a base solution (50 mM phosphate buffer, 13.5 mM KCl, 0.685 M NaCl and 10 mM MgCl_2_, 12.5% (v/v) glycerol, pH which was degassed by flowing Argon through buffers for at least 20 minutes before mixing or addition of buffers to samples. 1 mM paraquat dichloride and 1 mM ascorbic acid were added as reducing/oxidizing agents.^22^ Oxygen was depleted from the buffer by addition of 10 mM NaSO_3_,^29^ 50 nM protocatechuate-3,4-deoxygenase (*>*3 U/mg) and 2.5 mM of protocatechuic acid (ROXS PCD) or 0.66 M D-Glucose, 5000 U catalase, and 40-80 U glucose oxidase (ROXS GodCat). The ROXS GodCat buffer was additionally supplemented with 1 mM Tris(2-carboxyethyl)phosphine. All buffer components were obtained from Sigma Aldrich (Germany).

### Widefield fluorescence microscopy

If not stated otherwise, single-molecule fluorescence microscopy was performed on a custom-built inverted microscope (Nikon Eclipse Ti, Nikon, Japan) with epi-fluorescence and total internal reflection fluorescence (TIRF) illumination. The microscope setup included an autofocus system (PFS2) and a 100x 1.49 NA oil immersion objective (Apo TIRF, both Nikon). Images were recorded using a back-illuminated emCCD camera (iXon Ultra 897, Andor, UK) at 96 nm pixel size in the sample plane. A fiber-coupled multi laser engine (MLE-LFA, TOPTICA Photonics, Germany) equipped with 488, 561 and 640 nm laser lines was used for illumination. The excitation light was filtered by a quadband notch filter. A quad-band dichroic mirror separated the emission and excitation beam paths. Emitted signal was further filtered using 525/50 nm, 605/70 nm and 690/70 nm bandpass filters (all AHF Analysetechnik, Germany) mounted in a motorized filter wheel (FW102C, Thorlabs, US) placed before the camera. All microscope components were controlled using *µ*Manager.^47^ Exposure times, electron multiplying gain and illumination intensities were optimized for each sample individually to ensure maximum signals at the start of measurements while avoiding saturation of individual pixels.

### Widefield fluorescence microscopy with extended field of view and homogeneous illumination

Single-molecule trace acquisition with improved throughput was performed on a custom-built widefield fluorescence microscope built around an inverted Axiovert 200 stand (Zeiss, Germany). A 647 nm fibre laser with Gaussian-shaped emission profile (MPB Communications, Canada) was expanded to 6.0 mm and converted into a flat-top beam using a beamshaper (*π*Shaper, AdlOptica Optical Systems GmbH, Germany) and further expanded to a diameter of 47 mm. The expanded beam was then guided into the microscope stand and reflected towards a 100x 1.49 NA oil immersion objective (Apo TIRF, Nikon, Japan) objective. The variation in illumination power density was *<*15% across the entire illuminated field. Emitted signal was collected through the same objective, separated from excitation light using a quad-band dichroic filter (R405/488/561/635 Semrock, US) and further filtered using a 405/488/532/635 nm notch filter (Semrock) and a 700/50 nm bandpass filter (Chroma, US). Images were projected onto a back-illuminated sCMOS camera (Prime95B, Photometrics, UK). Samples were placed on a motorized stage (MS2000) and kept in focus using an auto-focus system (CRISP, both Applied Scientific Instrumentation, US). Camera and laser were synchronized using an Arduino Mega microcontroller board. All microscope components were controlled using *µ*Manager.

### Fluorophore stability measurements

The photostability of different fluorophores and the influence of ROXS buffers on the blinking and photostability of fluorophores was evaluated by recording time-lapse data from samples labeled with the corresponding fluorophore upon high intensity excitation. The stability of ATTO 647N was evaluated using DNA oligonucleotides labeled with ATTO 647N immobilized as described above. TMR and SiR substrates for SNAP-tag and HaloTag were evaluated in fixed cells using cell lines expressing NUP107-SNAP-tag or GlnA-HaloTag as described above. The stability of EGFP, mCherry and mNeonGreen was evaluated in COS-7 cells transiently expressing H2A-EGFP-HaloTag (kind gift of Richard Wombacher), TOMM20-mCherry-HaloTag^48^ or TOMM20-mNeonGreen (Allele Biotechnology, US) fixed 24 hours after transfection. For each fluorophore, the stability in PBS pH 7.4 and the NaSO_3_, ROXS PCD and ROXS GodCat buffer systems was tested with buffer compositions as described above. Prior to imaging, samples were washed once with PBS pH 7.4 and sample chambers were filled with the respective buffer and sealed with Parafilm to minimize gas exchange during experiments.

Bleaching curves were acquired on an epifluorescence setup for all buffer – fluorophore combinations described above. EGFP and mNeonGreen labeled structure were bleached at 577 W/cm2 average illumination power density, mCherry and TMR labeled samples were bleached at 839 W/cm2 average illumination power density and ATTO 647N and SiR labeled samples were bleached at 2.42 kW/cm2 average illumination power density. Image series were acquired with constant illumination until samples were fully bleached and the signal reached a plateau.

All data was background corrected by subtracting a constant offset from acquired image series to account for excitation light bleedthrough and sample autofluorescence. Offsets were manually determined for each sample and were well reproducible within one condition, but varied between conditions. Bleach curves were then extracted from image series by extracting the average intensity in each image within a masked region. Masks were obtained from a Gaussian-filtered average projection of the first 10 images in the series and local thresholding following the Bernsen method. Mask segmentation and intensity extraction was performed using custom-written code in Fiji/ImageJ. Bleach curves were then normalized by the maximum intensity in the respective trace and the half bleach time (*t*_1*/*2_) defined as the time at which the intensity traces had decayed to *<*50% of the maximum intensity was extracted using custom-written Matlab code.

### Photobleaching step analysis

The first 5 images from the measured image sequence were averaged and used to locate fluorophore clusters. The localization was performed with Fiji 1.52p ^49^,50 using the plugin thunderSTORM.^51^ Trace extraction was done with the trace extraction submodule of the quickPBSA package, as detailed in the package documentation. In short, the average signal from circular regions around the localizations was extracted with typical diameters of 950 nm for the in vitro samples and 150 nm for the NUP107 experiment. For background correction, the average signal from ring-shaped regions was subtracted (inner diameter 1.7 µm for origami, 0.6 µm for NUP107, outer diameter 2.0 µm for origami, 0.9 µm for NUP107). Regions around neighbouring localizations were excluded from the background region. Additionally, ROIs with nearest neighbours at a distance below 950 nm for DNA origami and 475 nm for NUP107 were excluded.

Photobleaching step analysis was performed using the quickPBSA package, as detailed in the main text and in the documentation of the quickPBSA package. Typically the threshold parameter for preliminary step detection was set at 0.03 and maxiter at 200. Other analysis parameters were kept at their default values, except for the mult threshold parameter in step refinement, which was typically set to 1.6 to decrease runtime.

Semi-synthetic datasets were generated by manual annotation of traces obtained from tetraATTO 647N DNA oligonucleotides measured in NaSO_3_ ROXS buffer. All analyses were carried out on a workstation with an 8-core CPU @3.4 GHz (Intel(R) Core(TM) i7-3770) and 12 GB DDR3 memory.

### Code availability

The quickPBSA package, example data and documentation are available under: https://github.com/JohnDieSchere/quickpbsa.

## Acknowledgement

We thank J. Shepard Bryan IV and Steve Pressé for fruitful discussions and Arina Rybina for critical reading of the manuscript. We also acknowledge funding from Deutsche Forschungsgemeinschaft (DFG) through project PhotoQuant HE4559/6-1.

