## Supplemental Information for "Photobleaching step analysis for robust determination of protein complex stoichiometries"

### Supplementary Information - Improved photobleaching step analysis for robust determination of protein complex stoichiometries

Johan Hummert,[†,‡,¶,‖] Klaus Yserentant,[†,‡,¶,§,‖] Theresa Fink,[†] Jonas Euchner,[†,‡,¶] and Dirk-Peter Herten[*,†,‡,¶]

†*Institute of Physical Chemistry, Heidelberg University, Heidelberg, Germany*

‡*Centre of Membrane Proteins and Receptors (COMPARE), Universities of Birmingham and Nottingham, UK*

¶*College of Medical and Dental Sciences & School of Chemistry, University of Birmingham, Birmingham, UK*

§*Faculty of Biosciences, Heidelberg University, Heidelberg, Germany*

‖*These authors contributed equally to this work.*

#### Fluorophore stability screen

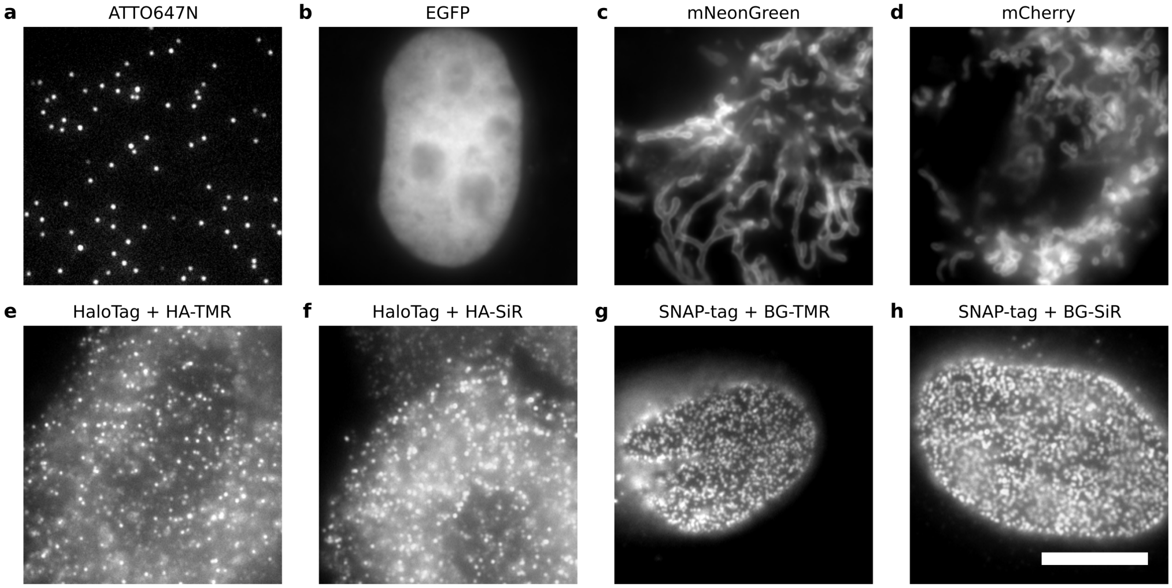


Figure S 1: Representative images from samples used for photostability characterization. **a,** DNA-conjugated and surface-immobilized ATTO 647N. **b,** COS-7 cells transiently expressing H2A-EGFP-HaloTag. **c,** COS-7 cells transiently expressing mNeonGreen-TOMM20. **d,** COS-7 cells transiently expressing TOMM20-mCherry-HaloTag. **e,f,** HeLa cells stably expressing GlnA-HaloTag. Cells were labeled with 100 nM HTL-TMR (e) or HTL-SiR (f) for 120 min. **g,h,** U2OS cells stably expressing NUP107-SNAP-tag were labeled with 200 nM BG-TMR (g) or BG-SiR (h) for 120 min. Images are representative for 7-10 cells from 2 independent experiments per conditions. Scale bar: 10 $\mu$m.

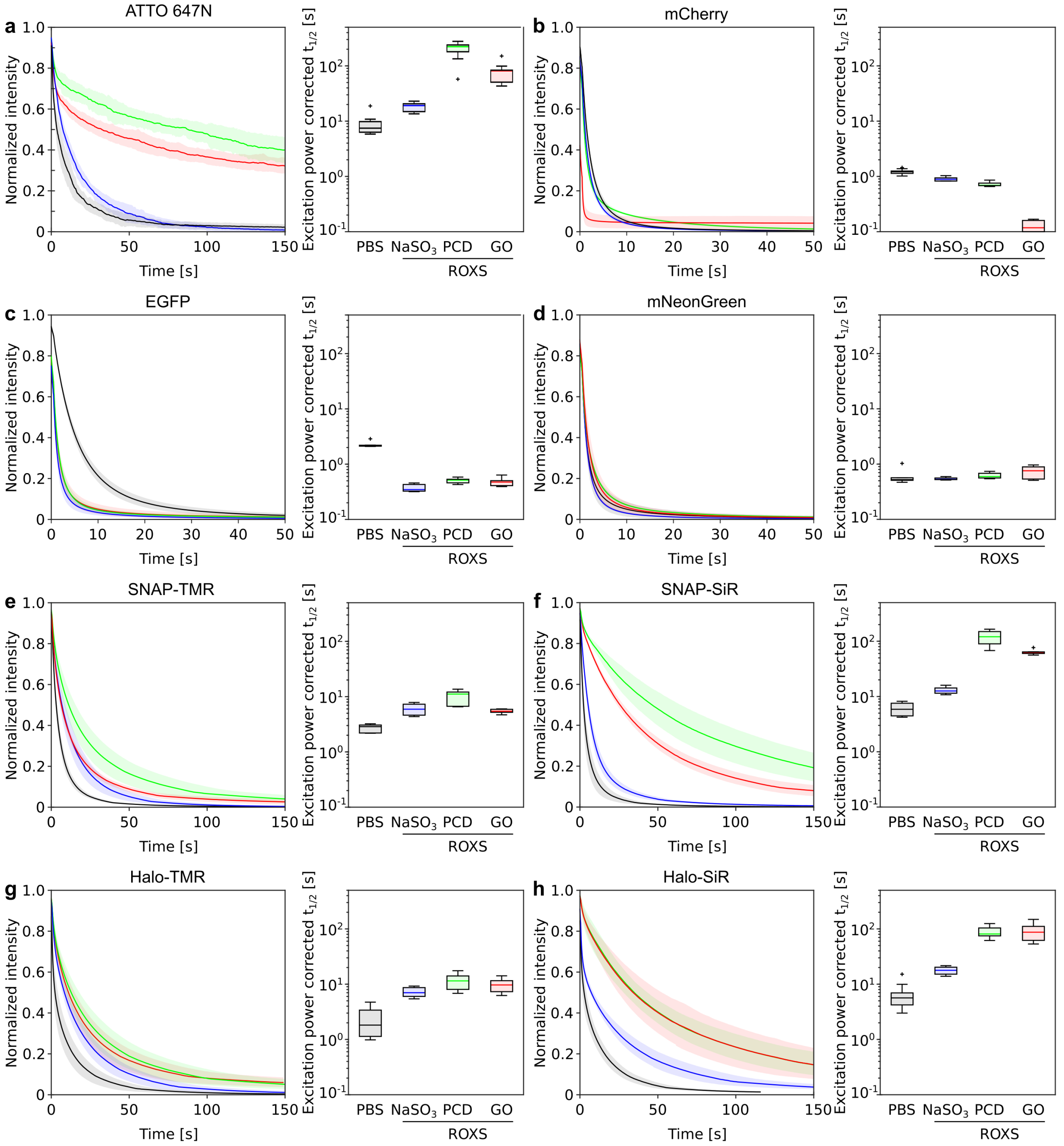


Figure S2: Photostability measurements for fluorescent proteins and protein tag substrates. Each fluorophore was exposed to high intensity illumination in respective buffers. Left: Photobleaching curves upon illumination for different buffers. Colors as in box plots. Mean (line) $\pm 1$ SD (shaded region) of 7-10 measurements from 2 independent experiments per condition. Right: Corresponding $t_{1/2}$ corrected for excitation power density. Box plots indicate median, 25$^{th}$ and 75$^{th}$ percentiles. Whiskers extend to 1x interquartile range, outliers are plotted as crosses.

Table S1: Spectroscopic properties of fluorophores used. Properties determined in PBS, pH 7.4-7.5 unless stated otherwise. $\epsilon$ – extinction coefficient, $\Phi_{Fl}$ fluorescence quantum yield, $\lambda_{peak}$ peak absorption wavelength, $\lambda_{ex}$ excitation wavelength. Spectral correction factors ($CF_{\lambda}$) were determined from published spectra or spectra measured for unconjugated BG/HA-dyes.

| Fluorophore | $\epsilon$ [x10$^3$M$^{-1}$cm$^{-1}$] | $\Phi_{Fl}$ | $\lambda_{peak}$ [nm] | $\lambda_{ex}$ [nm] | $CF_{\lambda}$ | Primary ref. |
|---|---|---|---|---|---|---|
| EGFP | 55.9 | 0.6 | 488 | 488 | 0.62 | Cormack et al. (1996)[1] |
| mNeonGreen | 116 | 0.8 | 506 | 488 | 1.00 | Shaner et al. (2013)[2] |
| mCherry | 72 | 0.22 | 587 | 561 | 0.64 | Shaner et al. (2004)[3] |
| BG-TMR | $89^{a,f}$ | $0.39^{b,g}$ | $555^{f}$ | 561 | 0.86 | Keppler et al. (2003)[4] |
| BG-SiR | $43.2^{c,g}$ | $0.30^{b,g}$ | $650^{g}$ | 640 | 0.75 | Lukinavicius et al. (2013)[5] |
| HTL-TMR | $78^{d,f}$ | $0.41^{d,f}$ | $548^{f}$ | 561 | 0.89 | Los et al. (2008)[6] |
| HTL-SiR | $130.2^{c,g}$ | $0.39^{b,g}$ | $648^{g}$ | 640 | 0.71 | Lukinavicius et al. (2013)[5] |
| ATTO 647N | $150^{e}$ | $0.65^{e}$ | 646 | 640 | 0.92 | ATTO-TEC |

[a] from Keppler et al. (2006)[7]
[b] from Lukinavicius et al. (2013)[5]
[c] from Erdmann et al. (2019)[8]
[d] Tetramethylrhodamine, from Grimm et al. (2015)[9]
[e] From ATTO-TEC specification sheet available at https://www.atto-tec.com
[f] Free dye
[g] Protein conjugate

#### Algorithm and benchmarking

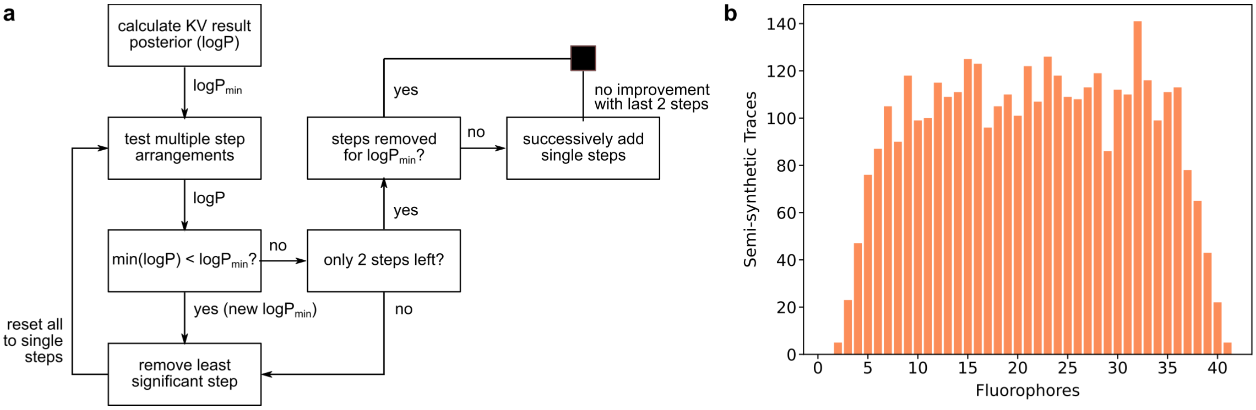


Figure S3: Algorithm and Benchmarking. **a,** Flowchart of the step refinement procedure to minimize the negative logarithm of the posterios (eq. 2) detailed in the manuscript. Additional breaking points are the maximum occupancy for a single step (max_mult) and the maximum number of added steps (max_added). **b,** Histogram of the number of synthetic traces included in the benchmark for each ground-truth fluorophore number.

The preliminary step detection (part II in the framework, detailed in the main text) relies on the Algorithm developed by Kalafut and Vischer,[10] which uses the Schwarz Information Criterion (SIC) to evaluate steps. In this context the SIC is defined as:

$$SIC(j_1, ..., j_k) = (k+2)log(n) + n \ln \hat{\sigma}^2_{j_1,...,j_k} + n \ln 2\pi + n \tag{1}$$

Steps are successively added until the addition of the step does not improve the SIC further. In the modified version used here steps where the difference in means, i.e. the mean signal before and after the added step, is below a defined threshold parameter, are not added even if they improve the SIC.

The step refinement (part IV in the framework, detailed in the main text) relies on the

Table S2: Symbols used

| SIC | |
|---|---|
| $j_1, ..., j_k$ | Step positions |
| $n$ | Timepoints in the trace |
| $\hat{\sigma}^2$ | sum over the variances of data between steps |
| **Posterior** | |
| $\theta$ | Bayesian parameters |
| $D$ | Data (photobleaching trace) |
| $K$ | Number of steps |
| $n_\phi$ | Timepoints in interval $\phi$ |
| $\sigma_f$ | Standard deviation of the single fluorophore signal |
| $\sigma_{bg}$ | Standard deviation of the background signal |
| $x$ | Signal at timepoint $l$ |
| $i$ | Active number of fluorophores at timepoint |
| $\mu_f$ | Mean signal of a single fluorophore |
| $\mu_{bg}$ | Mean background signal |
| $\lambda$ | Poisson distribution parameter for event occurences |
| $m$ | number of events (single fluorophore switches) |
| $d_y$ | number of data points between steps |
| $\gamma_0$ | Cutoff for the hyperparameter constraining $K$ to $m$ |

posterior from Pressé et al.:[11]

$$\begin{aligned}
-2\ln P(\theta|D) =& \sum_{\phi=0}^{K}\left(n_\phi \ln(i\sigma_f^2+\sigma_b^2) + \sum_{l=1}^{n_\phi}\frac{x_l - i\mu_f - \mu_{bg}}{i\sigma_f^2+\sigma_b^2}\right)\\
&+2\Bigg(-K\ln\lambda - \ln((m-K)!) - \ln K! - \ln(m-1)! + \sum_{y=0}^{m-K}\ln d_y!\Bigg)\\
&+2\Bigg(\gamma_0\frac{m-K+1}{K} + \ln(m-K+2) + \ln(m-K+1)\\
&\qquad - \ln(m-K+2) - (m-K+1)e^{-\gamma_0/K}\Bigg)
\end{aligned} \tag{2}$$

This expression is minimized iteratively, with fixed hyperparameters, using the algorithm in the flowchart (figure 3a).

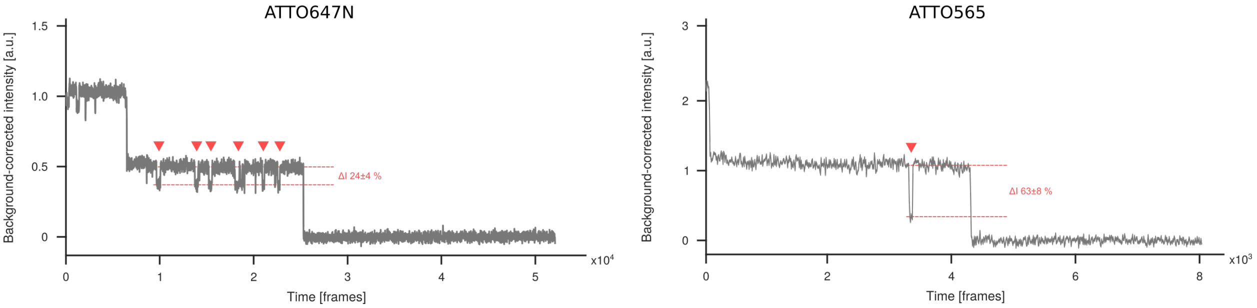


Figure S4: Representative photobleaching traces from the measurements with Atto 647N and Atto 565 illustrating the impact of fluorophore properties on photobleaching measurements. The second bright state of ATTO 647N is similar in brightness and does not affect the measurement strongly. In contrast the second bright state of ATTO 565 has a much lower brightness and could explain the overestimation observed in the *in vitro* measurement (figure 3 in the manuscript).

#### Counting of Nucleoporin 107

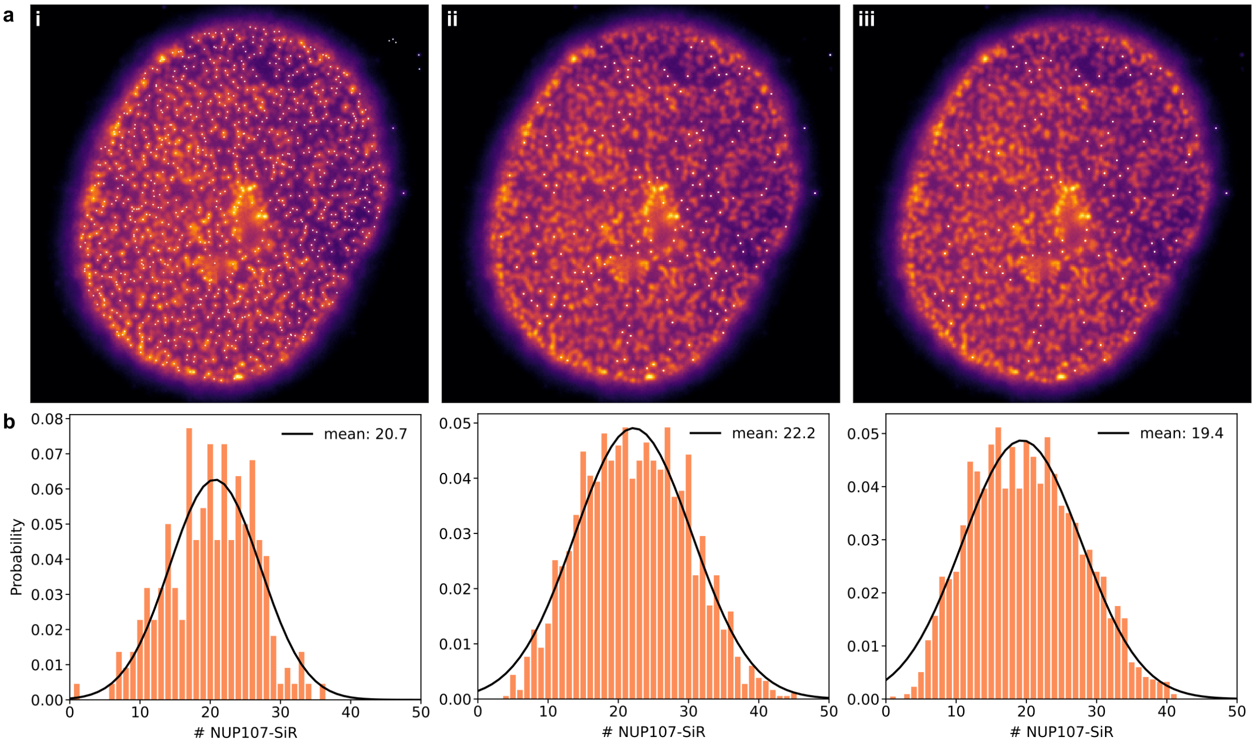


Figure S5: Counting of NUP107 **a,** Representative image with point localization and trace extraction. i) Initial ThunderSTORM localization result. ii) Remaining points after density filtering and filtering over gaussian width from localization. iii) Successfully evaluated spots (not flagged out, see quickpbsa package documentation). **b,** Extracted Fluorophore number distributions from the three independent measurements.
